# The U6 snRNA capping enzyme Amus/MePCE is required for the stability of U6atac and the processing of introns with features of both major and minor classes

**DOI:** 10.64898/2026.10.22.761030

**Authors:** Qiu Peng, Ying Xiao, Xiaolong Wu, ShaSha Wang, QianQian Liu, Juan Gong, Jiyong Liu, Min Cui, Zhiming Chen, Yikang S. Rong

## Abstract

Over 99% of the introns in humans are removed by the major spliceosome with the U6 snRNA as the catalytic component while the rest 1% by the minor spliceosome containing U6atac. Recent evidence supports mechanistic interactions between the two splicing pathways even though they have been considered as operating in parallel. Here we report a unique genetic condition in which both splicing pathways are partially impaired, and present evidence supporting inter-pathway interactions. We recently showed that the *Drosophila* Amus protein and its mammalian homolog MePCE serve as the methyltransferase of U6 and protect its stability. We show here that U6atac is greatly reduced under the same *amus* or *mepce* mutant backgrounds, suggesting Amus/MePCE serves the same protective function for U6atac. Interestingly, the processing of canonical minor introns is not significantly affected in the mutants, suggesting that a minimal amount of U6atac is sufficient for those reactions. In contrast, the splicing of a second class of “minor spliceosome sensitive” introns are inhibited. These introns harbor splice sites different from the minor consensus even though they interact favorably with the minor spliceosome as shown by others. Interestingly, most members of this class of minor-like introns are evolutionarily “younger” than their canonical counterparts, and our in-depth analysis of one such intron supports that some of them might be functionally neutral. Our study provides empirical evidence supporting the existence of a class of introns jointly regulated by the two splicing pathways, and revealing additional complexity in the functional evolution of minor splicing.

## Introduction

The reaction of pre-mRNA splicing, in which exons are joined with the intervening intronic sequences excised, is carried out by a mega-dalton protein and RNA complex called the Spliceosome. Central to the spliceosome are five U-rich small nuclear RNAs (snRNAs) with distinct functions: to define exon/intron boundaries (U1 and U2); to catalyze the chemical reaction in splicing (U6); to regulate the aforementioned catalytic activity (U4); and to serve as a scaffold for exon alignment (U5) (for reviews on spliceosomes and spliceosomal snRNAs see Matera and Wang 2014; and Wilkinson et al. 2020). The above-described Spliceosome, named the major spliceosome, processes over 99% of the introns in the human transcriptome (Sheth et al. 2006; Parada et al. 2014). The 1990s saw the discovery of another, minor, class of introns (Jackson 1991, Hall and Padgett 1994; Tarn and Steitz 1996), which are processed remarkably similarly to the major class, but using the minor spliceosome. In the minor spliceosome, four functionally similar snRNAs replace the ones in the major spliceosome (U11 for U1, U12 for U2, U4atac for U4, U6atac for U6), in addition to several protein factors (reviewed in Akinyi and Frilander 2021; Ding et al. 2023).

Central to both splicing machineries are the U6-type catalytic snRNAs: U6 and U6atac, the latter named after the dinucleotides (AT and AC) often present at the ends of a minor intron. Both U6 and U6atac are produced by RNA polymerase III (Pol III), while the other spliceosomal snRNAs are produced by Pol II. U6 is protected by a 5’ methylated phosphate cap (Shumyatsky et al. 1993). Based on their very similar functions, and essentially identical biogenesis, it is expected that the two snRNAs are protected similarly by the 5’ cap. In addition, a cryo-EM structural model is consistent with the presence of a 5’ methyl-phosphate cap on U6atac (Bai et al. 2021). However, whether U6atac is protected by the presumed 5’ cap lacks empirical evidence, and the biological consequences of missing the cap of U6atac remain uncharacterized. We previously showed that the *Drosophila* Amus methyltransferase and its mammalian homolog MePCE control 5’ capping of U6, so that even a reduced level of either Amus or MePCE causes a reduction in the steady state level of U6 in cells (Peng et al. 2023). Here we provide evidence supporting that Amus/MePCE fulfills an identical function for U6atac protection.

The two splicing pathways, major *versus* minor, have long been considered as operating in parallel despite their catalytic similarities and shared protein and RNA components (e.g. U5 snRNA). As the two chemical reactions are catalyzed by two different tri-snRNPs (U4:U5:U6 for major and U4atac:U5:U6atac for minor introns), it is conceptually possible that a less stringent assembly process generates a “hybrid” spliceosome capable of acting on the “opposite” class of introns, as has been demonstrated experimentally (Shukla and Padgett 2004; Dietrich et al. 2009). In addition, through a large scale bioinformatic analysis of introns from 263 genomes, unique classes of “minor-like” and “hybrid” introns are identified that share common features with both canonical minor introns and major ones (Olthof et al. 2024). In addition, upon functional perturbation of the minor spliceosome, many of those introns experience splicing inhibition suggesting that they are processed by the minor splicing machinery. Moreover, a genetic study in *Drosophila* using null mutants identifies a novel class of introns that are sensitive to the loss of U6atac or U12 snRNAs (Li et al. 2020). These “minor spliceosome sensitive introns” (minSIs) possess splicing elements with minor-like features, and likely represents a subclass of the “minor-like” introns identified via bioinformatics (Olthof et al. 2024). As these introns are different from the canonical minor introns, they have the potential to be processed by the major spliceosome as well (Li et al. 2020; Olthof et al. 2024).

In this study, we showed that *amus* mutations in *Drosophila* destabilize the catalytic snRNAs for both spliceosomes. We discovered that canonical minor introns are not affected by the partial loss of U6atac, while splicing of minSIs are. We argue that this is due to the concurrent partial loss of U6, a critical component of the major machinery. In addition, *Drosophila* minSIs are generally “younger” evolutionarily than canonical minor introns, consistent with them being in a transitory state as suggested for the “minor-like” introns (Olthof et al. 2024). Finally, we rule out RNA exonuclease Rai1 as the enzyme responsible for the degradation of unprotected U6-type snRNAs.

## Materials and Methods

### *Drosophila* stocks and crosses

All *Drosophila* stocks were obtained from the Bloomington Drosophila Stock Center, Indiana, USA, unless mentioned otherwise. The *Df(3R)BSC47* was used as a deficiency of the *amus* region (Bloomington stock #7443). The *rai1^e04371^* was used as a second allele of *rai1* (Bloomington stock #85169), in which a piggyBac transposon inserts into the second coding exon of *rai1*. The *w^1118^* stock is used as controls. The alleles of *amus^1-1^* (three amino acid changes) and *amus^A-1-3^* (10 bp deletion), transgenic lines of *amus^Y84A^* and *amus^GFP^*, and transgenic lines of *UAS*-driven *amus* cDNA and the cDNA for the Amus-MePCE hybrid protein, have been described in Peng et al. 2023. The *hiphop^HA^* allele has a 3X-HA tag inserted at the endogenous *hiphop* locus, which was described in Cui et al. 2021. The *hiphop^N-GFP^* line has a *hiphop* transgene with an N-terminal GFP tag, which was described in Chen et al. 2022. The *U6atac^Δ^* mutant was a kind gift from Dr. Yong-zhen Xu at Wuhan University, China. Stocks of *D. simulans*, *D. yakuba*, *D. pseudoobscura*, *D. willistoni* and *D. virilis* were kind gifts from Dr. Yirong Wang of Hunan University, China. The *bin3^2-7^* and *bin3^4-7^* alleles were kind gifts from Dr. Steven Hanes at SUNY Upstate Medical University, USA.

Transgenic stocks generated in this study include *UAS*-controlled *amus^Y83F^* and *amus^Δ76^* cDNAs based on the pUAST-attB vector. The transgenes were inserted into an *attP* landing site at 75 on chromosome *3L*. They were recombined onto the *amus^A-1-3^* chromosome *3R* by meiotic recombination. An *actin-gal4* transgene on chromosome *2* was used to drive their expression. The *rai1^2-4^* mutation was generated via CRISPR/Cas9 using the sgRNA with the sequence of 5’-ACACACCGATGCACTTCGGG**CGG**, with the PAM sequence in bold.

### RNA related experiments

Total RNA was extracted with a standard TRIzol reagent-based method. For Northern Blot analyses, 1μg of total RNA was denatured at 80°C for 3 min and separated on 5-8% 8M Urea-polyacrylamide gels in TBE buffer. Northern blotting was carried out by standard protocols and probed with biotin labelled oligonucleotides synthesized by Sangon Biotech China, and their sequences are listed in Table S1. Hybridization was performed overnight at 37°C for the U6atac and U4atac probes, whereas all other probes were hybridized overnight at 42°C. The signals were generated with streptavidin conjugated HRP, and quantified by Image lab software 5.2 of Biorad.

For RT-PCR, total RNA was extracted from 10 to 15 adult flies or third instar larvae using TRIzol reagent. RT was carried out with the PrimeScript Kit from TaKaRa. Primer sequences are listed in Table S1.

Human HEK293T cells were treated with siRNA reagents as previously described (Peng et al. 2023). An anti-MePCE antibody was purchased from Bethyl Laboratories Inc. (A304-185A), and used at a dilution of 1:1000 on Western blots.

### BLAST analyses of minor introns in *Drosophila* species

DNA sequences of the 5’ splice site (SS) and 3’SS of the minor and minS introns were taken from Supplemental Data 3 in Li et al. 2020, which contain 80nt at each splice site. A DNA fragment of 160bp (80+80) was used in BLAST searches from the Flybase server with the first twelve sequenced *Drosophila* species selected. A species is classified as containing a particular intron when both splice sites return homologous sequences that are separated by a bp distance comparable to the size of the original intron.

## Results

### Amus or MePCE partial loss of function destabilizes U6atac

We previously generated mutations to the *Drosophila amus* gene, which possesses a methyltransferase domain homologous to those in the *Drosophila* Bin3 and mammalian MePCE proteins (Peng et al. 2023). Null mutants of *amus* (*amus^A-1-3^*) die at the first to second instar larval stages accompanied by a greatly reduced level of the U6 snRNA in mutant cells. This is consistent with that Amus is the capping enzyme for U6 in *Drosophila*, and we showed that this role is carried out by MePCE for mammals (Peng et al. 2023). Interestingly, a hypomorphic mutation of *amus* (*amus^1-1^*) supports viability but not fertility, and we uncovered splicing defects in the form of intron retention in *amus^1-1/A-1-3^* testes (Peng et al. 2023), consistent with a reduced U6 level in *amus^1-1/A-1-3^* adults (Figure 1A). Remarkably in the same adults, U6atac is also reduced strongly (Figure 1A). This reduction is due to the loss of Amus function, as a transgene carrying the wildtype genomic locus of *amus* was able to restore both the U6 and U6atac levels (Figure 1A).

**Figure 1.**
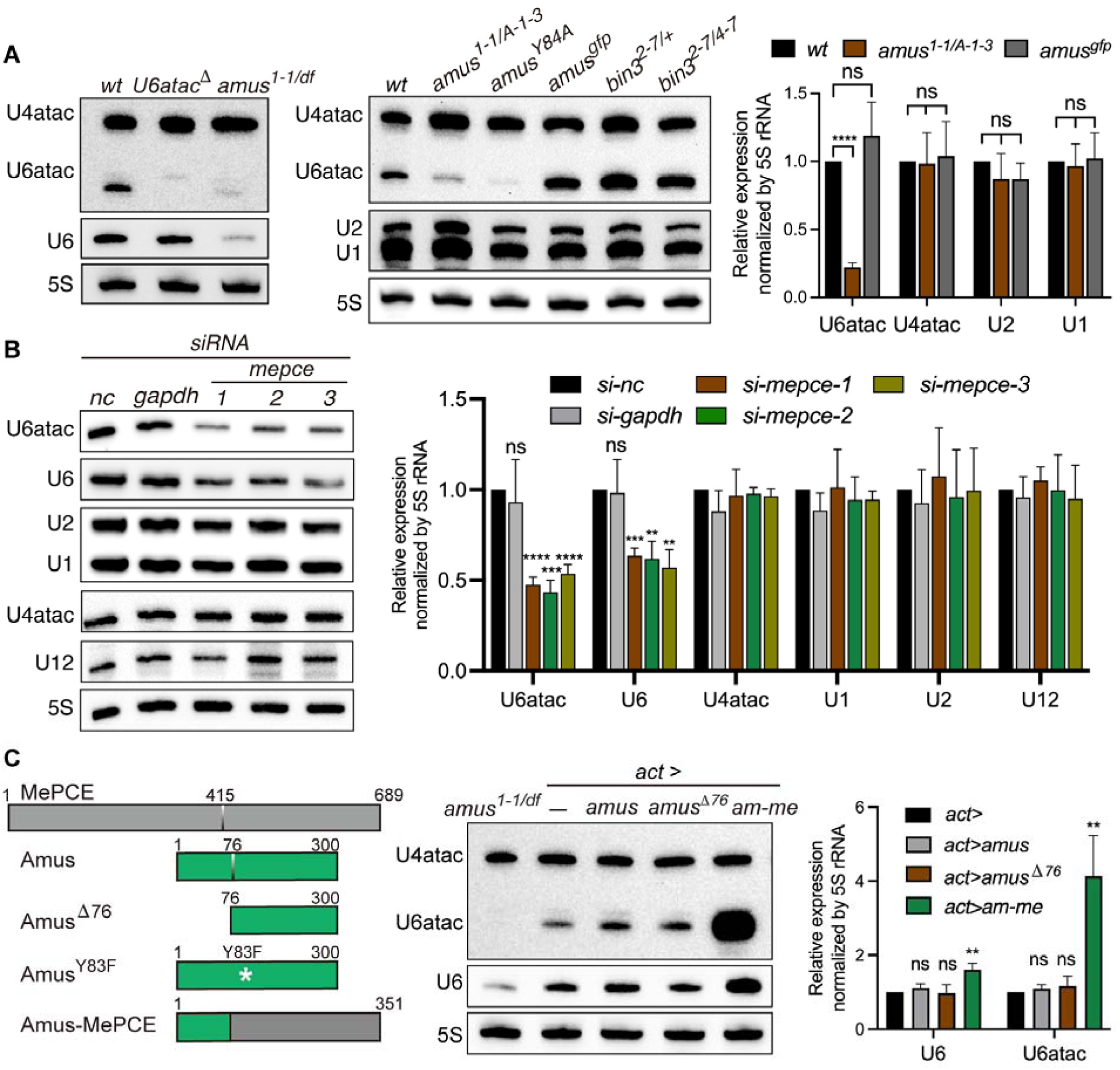
Amus/MePCE is essential for maintaining the level of U6atac. **A. Amus is essential for U6atac stability in *Drosophila***. In the two panels showing Northern blot analyses of small RNAs, genotypes are listed on top (*df*: a chromosomal deficiency of the *amus* region), and the small RNA species being tested are indicated to the left. In the middle panel, the full genotype of *amus^Y84A^* (another hypomorphic mutation of *amus*) is *[amus^Y84A^] amus^A-1-3^*, and that of *amus^GFP^* is *[amus^GFP^] amus^A-1-3^* (rescue of *amus* mutation). Loss of Bin3, the capping enzyme for *7SK* in *Drosophila*, does not affect U6atac. The right panel shows data quantification for the middle panel from three biological replicates. ns: not significant. ****: p<0.0001. **B. MePCE is essential for U6atac stability in human cells**. On top of the panel showing Northern blot results are RNAi treatment conditions. “nc” indicates a non-targeting siRNA as a negative control, with siRNA against *gapdh* as an additional control and three siRNA regents against different parts of *mepce* as previously described (Peng et al. 2023). For the efficacy of siRNA KD, see Figure S1A. The small RNAs of interest are listed at the left. The right panel shows data quantification of the Northern blots from three biological replicates. ns: not significant. **, p<0.01 ***: p<0.001, ****: p<0.0001. **C. The methyltransferase domain of MePCE can functionally replace that of Amus**. At the left are diagrams of different proteins introduced into the *amus* mutants. At the top is the 689 residues long human MePCE protein with a methyltransferase domain starting at residue 415. Underneath MePCE is the Amus protein with a methyltransferase domain starting at residue 76. The Tyr83 residue critical for methyltransferase activity is mutated to Phe in the Amus^Y83F^ mutant. At the very bottom is the Amus-MePCE chimeric protein (Am-Me). The middle panel shows Northern blots probing for various small RNAs, indicated to the left. The genotypes are listed at the top. The leftmost lane contains RNA from pupae of *amus^1-1/df^*. The second to left lane contains pupal RNA from animals with the *actin-gal4* transgene (act>), and heterozygous for the *amus^A-1-3^* mutation but possessing no *amus* cDNA transgene (-). The rest three samples were from animals with the *actin-gal4* transgene (act>), homozygous for *amus^A-1-3^*, and possessing: a cDNA for wildtype *amus* (amus); or a cDNA for Amus missing the first 76 residues (amus^Δ76^); or a cDNA for the Amus-MePCE chimeric protein (am-me). The Amus^Y83F^ expressing animals died very early in development and were not included in Northern Blot analysis. The right panel shows data quantification from three biological replicates. ns: not significant. **: p<0.01.

In human HEK293T cells that have been treated with *mepce* RNAi agents as previously described (Peng et al. 2023), we also observed greatly reduced U6 and U6atac levels upon an effective MePCE knockdown (Figure 1B and Figure S1A). In lieu of a *mepce* rescue experiment in human cells, we used a scheme previously designed to rescue defects in snRNA production in *Drosophila* cells by expressing the human enzyme (Peng et al. 2023). In this scheme, the MePCE methyltransferase domain was expressed in *Drosophila* as a fusion protein attached to the N-terminal 76 amino acid residues from Amus, which we named Am-Me (Figure 1C). The expression of Am-Me was accomplished by the Gal4-*UAS* system using an *actin5C*-driven *gal4* transgene for ubiquitous expression, and a *UAS* controlled *am-me* cDNA. As controls, we reintroduced into the *amus* null background three other cDNA constructs, (1) a wildtype copy of *amus*, (2) an *amus* carrying a point mutation disrupting the methyltransferase active site (Y83F), and (3) a truncated *amus* missing the first 76 amino acids. These four transgenes, along with *actin-gal4*, were individually introduced into the *amus^A-1-3^* null mutant background. All except the “enzyme-dead” *amus* mutant (*amus^Y83F^*) were able to rescue significant development to the mutants, with the wildtype gene supporting adult viability and the other two conditions (*am-me* and *amus^Δ76^*) supporting pupal development. Under the three “rescued” conditions, the levels of U6 and U6atac were fully restored, and sometimes even over and above the wildtype levels (Figure 1C). In particular, expression of the Am-Me protein led to a vastly elevated U6atac level. We used a serum previously raised against the full length Amus protein (Peng et al. 2023), in Western blotting with extracts from whole pupae. We detected an elevated level of Am-Me protein in comparison to that of either the wildtype or the N-terminally truncated Amus proteins (Figure S1B). We note that results from this Western blot analysis likely underestimate the overproduction of Am-Me as the chimeric protein lacks over two-thirds of the potentially antigenic region in Amus. Therefore, we suggest that the level of U6atac is positively affected by the level of the methyltransferase that caps it. Hence, we showed that mammalian methyltransferase is able to protect U6atac in flies and it is reasonable to suggest that it functions similarly in human cells.

### U6atac is in large access for canonical minor splicing

In light of the greatly reduced U6atac level, even under an *amus* hypomorphic background, we suspect that the splicing of minor introns is impacted in the *amus* mutants. We first focused on the 15 introns in *Drosophila* that have been classified as canonical minor introns exclusively processed by the minor spliceosome (Li et al. 2020; Olthof et al. 2024). We used an RT-PCR based assay in which a pair of exon-located primers amplify a fragment from cDNA species with or without the intron of interest (Figure 2A). To our surprise, none of the 15 canonical minor introns show discernible retention in *amus^1-1/A-1-3^* adults (Figure 2C). On the other hand, using a previously characterized complete deletion of the *U6atac* locus (*U6atac^Δ^*), we were able to recapitulate the splicing defects in 14/15 of these introns in the homozygotes (Figure 2B). We were left to wonder whether the residual amount of U6atac in the *amus* mutants is sufficient to support the processing of these introns. To further reduce U6atac, we introduced a *U6atac^Δ^* heterozygous mutation into the *amus^1-1/A-1-3^* background (full genotype: *U6atac^Δ/+^; amus^1-1/A-1-3^*). Remarkably, we were then able to detect splicing defects in 10/15 of the canonical minor introns (Figure 2C). These results suggest that the function of the minor spliceosome must not have been critically impaired under the *amus^1-1/A-1-3^* background, despite the appearance of a severe loss of U6atac RNAs on Northern blots. This situation is reminiscent of a previous study in which a less than expected defects in minor splicing was observed in another *U6atac* allele generated by the insertion of a *P* element (Pessa et al. 2010). It was suggested that the *P* element allele retains substantial U6atac function even though animals homozygous for the mutation fail to survive. Therefore, we conclude that partial loss of Amus function creates a sensitized background in which the minor splicing function is compromised though not abolished.

**Figure 2.**
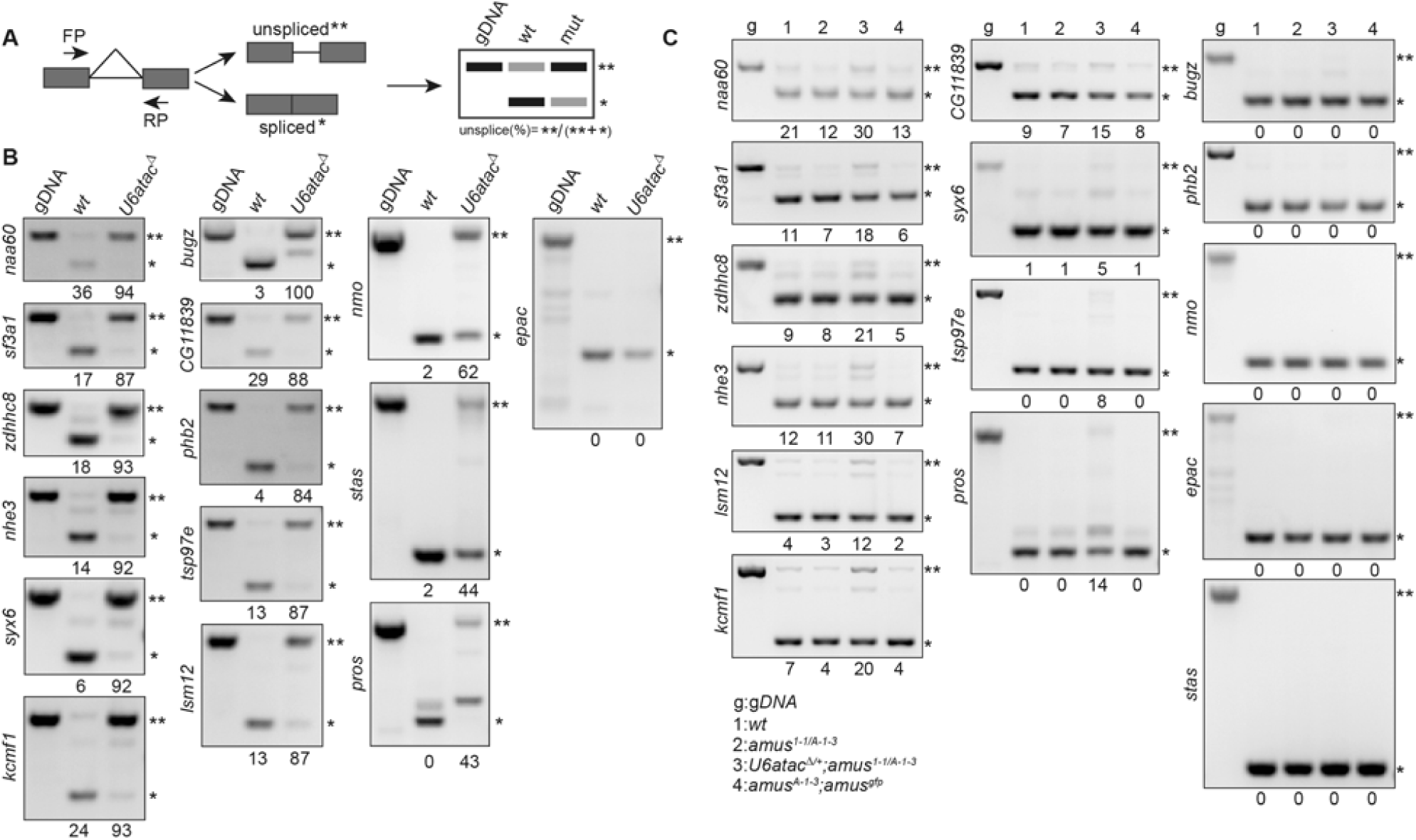
Splicing of canonical minor introns is insensitive to the partial loss of Amus. **A. The RT-PCR based assay for measuring splicing efficiency.** A hypothetic RT-PCR is depicted to the left in which a pair of primers (FP and RP) derived from exons (rectangular boxes) are used to amplify cDNA. Two possible products are produced: the “unspliced” (**) product that includes the intron (horizontal line) and the “spliced” (*) product. At the right is a hypothetic DNA gel picture when the PCR products are run. As a control, genomic DNA (gDNA) was used as a PCR template to mark the running position of the unspliced product. The genotypes are listed above, and the two products are labelled. Underneath the picture, quantification of the relative abundance of the two products are given as the percentage of the “unspliced” product over the total products. **B. The splicing of minor introns depends on U6atac.** The splicing of all except one previously defined canonical minor introns are inhibited by a deletion mutation of *U6atac*. Total larval RNA was used for cDNA synthesis. The names of the gene where the intron of interest resides are listed to the left of the gel pictures. C. The effect of *amus* mutation on the splicing of canonical minor introns. RNA used for cDNA synthesis was isolated from adults with the genotype indicated on top of the gel pictures, and listed at the very bottom. g: genomic DNA as the template of PCR.

### The processing of “minor spliceosome sensitive introns” are sensitive to the loss of Amus

In a prior study of minor splicing in *Drosophila*, Li et al. (2020) identified one additional class of introns, whose processing is inhibited upon the complete loss of the minor splicing machinery (deletion of either *U6atac* or *U12* genes), and termed those as minor spliceosome sensitive introns (minSIs). These introns do not process all of the consensus features of a canonical minor intron, nevertheless the majority of them (35 out of 48) possess a 5’ splice site (5’SS) capable of pairing with *U11* snRNA with a favorably small free energy (Figure 5A in Li et al. 2020), and some of them were able to associate with minor spliceosome specific protein components (Li et al. 2020).

Using the RT-PCR based assay shown in Figure 2A, and the *U6atac^Δ^* null mutation, we were able to demonstrate splicing defects in at least 34 of these 48 minSIs (Figures 3A, S2A, and summarized in Table S2). In contrast to the 15 canonical minor introns that are essentially irresponsive to Amus’s partial loss of function, we detected splicing inhibition in 24 out of the 34 RT-PCR validated minSIs (Figure 3B and Figure S2B). Aided by a chromosomal deficiency of the *amus* locus (*df*), we constructed another Amus partial loss of function background (genotype *amus ^1-1/df^*), and again observed splicing defects in minSIs but not canonical minor introns (Figure S2D). To establish that the effect of the *amus* mutation is specific to minSIs, but not minSI-containing genes, we randomly picked five of the 24 minSI-containing genes, and performed the same RT-PCR splicing assay on a major intron. As shown in Figure S2C, none shows defects in splicing. We also tested the effect of further reducing the dosage of *U6atac* on minSI splicing, using the heterozygous deletion mutation as previously done (Figure 2C and Figure S2B). In general, we did not observe a worsening effect on minSI splicing when the function of minor splicing was further impaired (Figure 3B). We visually inspected whether these 24 affected minSIs are ones tending to have 5’SS well paired with *U11*, but did not observe a significant correlation. While 18/24 affected minSIs were classified as “well paired”, 8/10 unaffected minSIs were as well (p=0.7541 from a Chi-square test).

**Figure 3.**
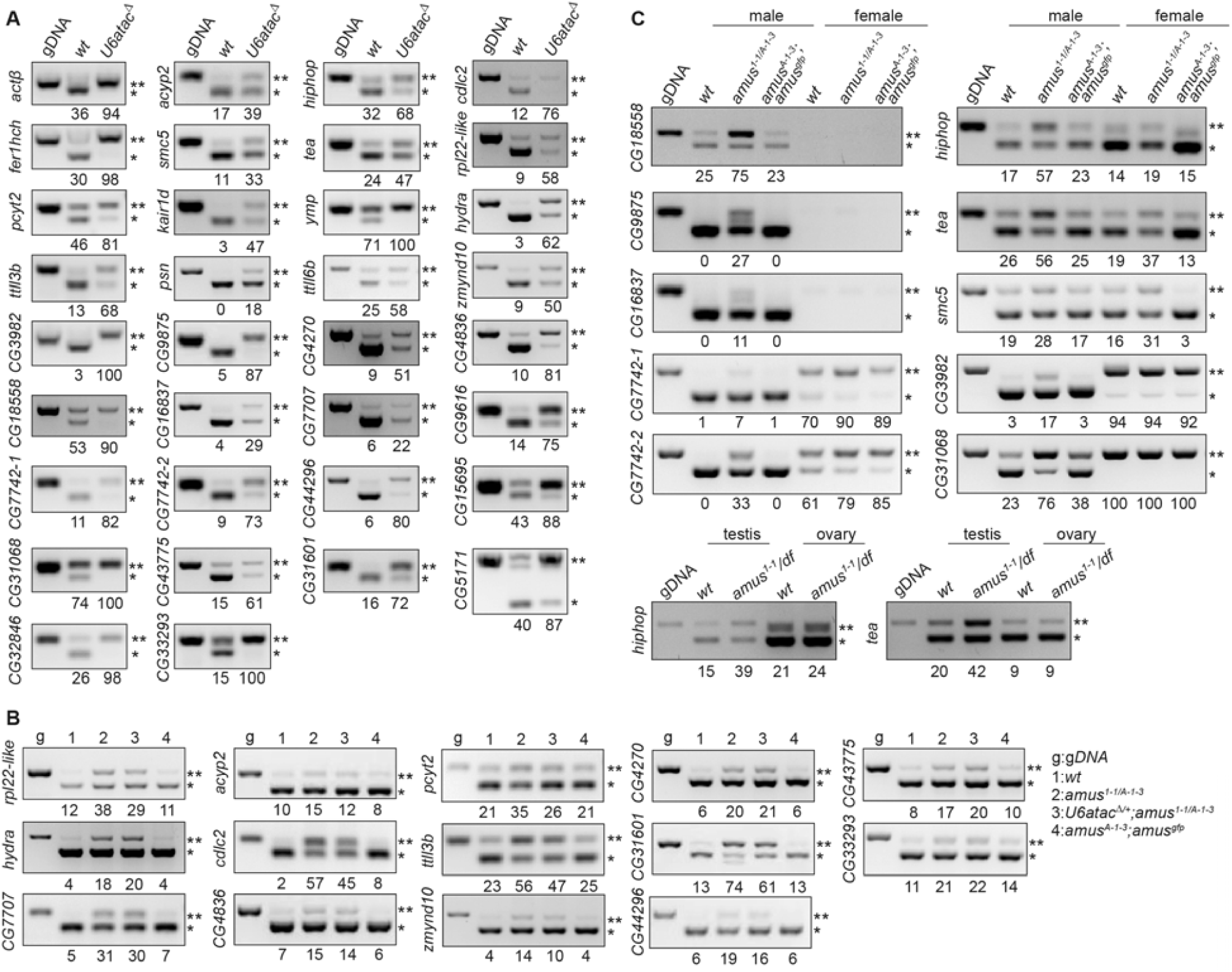
Splicing of minSI requires full function of Amus. **A. minSIs rely on U6atac for normal processing.** Larval RNA was isolated for the RT-PCR based assay described in Figure 2A. The processing of 34 out of 48 minSIs were defective. **B. The effect of *amus* mutations on minSI splicing**. RNA used for cDNA synthesis was isolated from adults with the genotype indicated on top of the gel pictures and listed at the very bottom right. g: genomic DNA was used in PCR. **C. Sex specific effects of *amus* on minSI splicing.** RNA was isolated from males and females of the three genotypes (wildtype, *amus* mutant and *amus^GFP^* rescue). The left panel shows five examples of male enriched expression and the effect from *amus* mutations. The right panel shows examples in which the effect from *amus* mutation appears greater in males than in females, such as the minSIs in *hiphop* and *tea* genes. The bottom panel shows splicing assay results using RNA isolated from dissected testes and ovaries of *wt* and mutant animals.

Interestingly, some of the 24 affected introns either have a sex biased expression, or the effect from the *amus* mutation was more pronounced in male adults (Figure 3C). Using dissected ovaries and testes, we showed that the male-biased effect might be largely originated from germline tissues (Figure 3C), at least for the two genes (*hiphop* and *tea*) that encode proteins essential for telomere protection (Gao et al. 2010; Zhang et al. 2016).

Therefore, minSIs are more sensitive to the partial loss of U6atac in the *amus* mutants than canonical minor introns, perhaps due to the fact that the *amus* mutation also affects the major spliceosome through the destabilization of U6.

### Minor spliceosome sensitive introns in *Drosophila* are generally “young”, and some might be functionally neutral

A recent bioinformatic analysis of exon/intron structures came to several interesting conclusions (Olthof et al. 2024). First of all, the binary classification of introns to major or minor classes does not reflect the full spectrum of sequence diversity of splicing elements. Rather, there are “minor-like” and “hybrid” introns that retain to various degrees of the features defined for canonical minor and major introns. In addition, a significant portion of these introns resemble the minSIs, which is the interest of this work, in that their processing is inhibited upon perturbation of the minor spliceosome. Lastly, these introns have been suggested to be in a transitory state evolutionarily, converting from a minor to a major intron and *vice versa*. We therefore characterized the minSIs that we have studied for their evolutionary trajectory.

Simple BLAST searches using flanking DNA sequences from the 5’ and 3’ SS of the introns identify the state of a particular intron in a *Drosophila* species as either present or absent. We focused on the twelve initially sequenced *Drosophila* species (Crosby et al. 2007). While three of the 15 canonical minor introns (shown in red in Figure 4A) are limited to species of the *melanogaster* subgroup, the rest are present in more diverged species, including >50% (8/15) being present in the Hawaiian *Drosophila* of *D. grimshawi*, with over 40 million years separating it from *D. melanogaster* (Russo et al. 1995). In contrast, only one of the 24 minSIs (shown in blue in Figure 4A), which responded to the loss of Amus function, is present outside of the *melanogaster* subgroup. In addition, 10 of 24 are limited to *D. melanogaster* and its closest relatives of *D. simulans* and *D. sechellia*. This result is consistent with the notion that canonical minor introns are generally more conserved than minor-like introns (Olthof et al. 2024). Therefore, we suggest that the minSIs in *Drosophila* are in general evolutionarily younger, consistent with the proposal they are in a transitory state.

**Figure 4.**
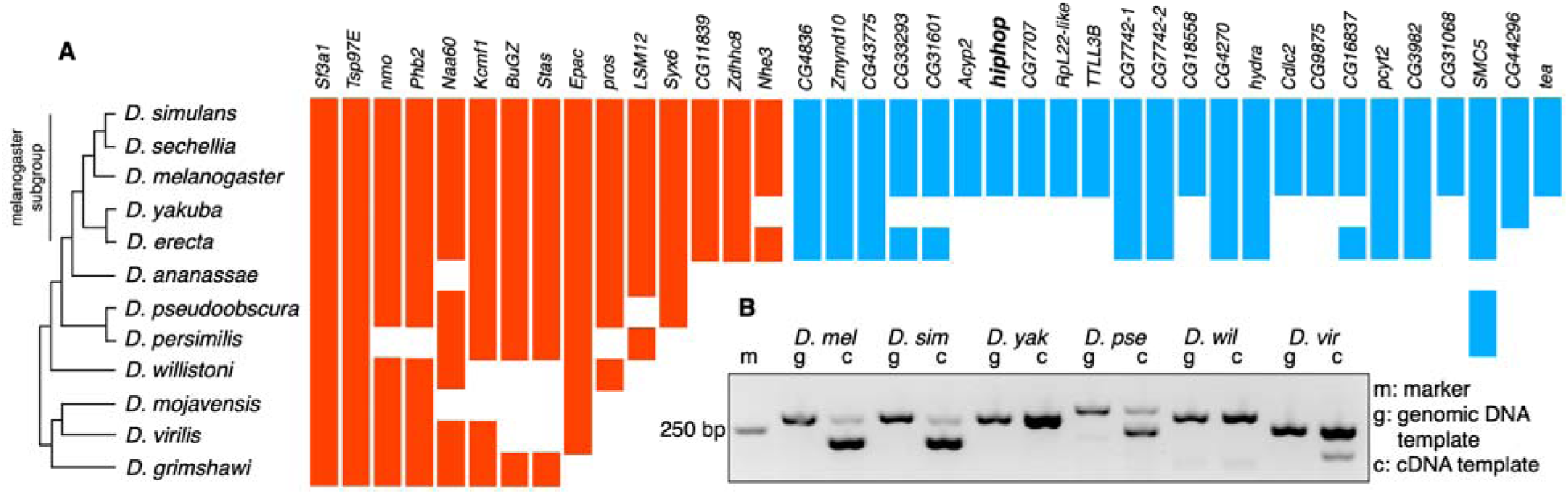
minSIs in *Drosophila* can have dynamic evolution. **A. The presence or absence of canonical minor introns and minSIs in *Drosophila* genomes.** Shown to the left is a cladogram illustrating the order of the different species. To the right of the cladogram, each intron (listed on top) is classified as present (filled space) or absent (empty space) based on BLAST searches for homologous splice sites. Canonical minor introns are colored in red and minSIs that are affected by the *amus* mutation in blue. **B. The splicing of the intron in *hiphop* genes from different *Drosophila* species.** RT-PCR was performed with species specific primers using RNA from adults of the following species: *D. mel*, melanogaster; *D. sim*, simulans; *D. yak*, yakuba; *D. pse*, pseudoobscura; *D. wil*, willistoni; and *D. vir*, virilis. A control PCR with genomic DNA as template was used to indicate the running position of the unspliced product. See Figure S3 for the precise intron positions in *D. mel*, *D. sim*, *D. pse* and *D. vir*.

To provide empirical data to support our genomic analyses, we characterized the minSI in the *hiphop* gene, which we had studied (Gao et al. 2010; Cui et al. 2021; Chen et al. 2022). The *hiphop* gene in *melanogaster* has a minSI in the 5’UTR with a 3’SS of AG^ATG in which ATG is the START codon. Using total RNA isolated from adults, we investigated the presence of a similar intron in *D. simulans*, *D. yakuba*, *D. pseudoobscura*, *D. willistoni*, and *D. virilis*. Consistent with our BLAST results, a minSI at the expected position is efficiently spliced in *D. simulans*, but not in *D. yakuba*, nor in the more distantly related *D. willistoni* (Figure 4B). Remarkably, RT-PCR detected splicing of an intron of the expected size and at the expected position from the *hiphop* genes of *D. pseudoobscura* and *D. virilis* (Figure 4B). Sequencing of the PCR product confirms its existence, and interestingly, the 3’SS of AG^ATG also contains the START codon of the HipHop protein (Figure S3). These results suggest the existence of a more ancient intron at a common position. One likely scenario is that this intron is lost in some *Drosophila* lineages but converted to a minSI in the common ancestor of *melanogaster* and *simulans*. During our studies of *hiphop*, we have generated two *hiphop* alleles in which the insertion of N-terminal tags just downstream of the ATG codon inevitably introduced diverse sequences to the 3’ SS of the minSI. The wildtype 3’ SS with the sequence of 5’-tattccAG^ATGgcctcca is changed to 5’-tattccAG^ATGtacccat in the *hiphop^HA^* allele, and to 5’-tattccAG^ATGagcaagg in the *hiphop^N-GFP^* allele. Remarkably, the splicing of the minSI is not affected by the sequence alteration to the immediate vicinity of the 3’SS in these new *hiphop* alleles (Figure S4A), which suggests that the signal for the elimination of the minSI is primarily embedded in the intron itself.

As this intron lies in the 5’UTR of *hiphop*, its biological function, if any, would likely be for regulating the production of the HipHop protein, either through transcription or more likely translation. Loss of HipHop leads to telomere uncapping and telomere fusion, which can be observed from mitotic squashes on larval neuroblasts (Gao et al. 2010; Cui et al. 2021). In *U6atac^Δ^* mutants where we observed the strongest defect of minSI splicing in *hiphop*, we scored 30 mitotic figures in 4 mutants but did not observe any chromosome configuration suggestive of telomere fusions. The splicing of the minSI in *hiphop* is also affected by the *amus* mutation. However, the fact that *amus* mutants survive as normal looking adults is indicative of the absence of telomere uncapping, which is expected to be lethal. We previously showed that HipHop is also required for the suppression of the telomeric transposons in the adults (Cui et al. 2021). As shown in Figure S4B, the expression of these elements is not affected by the *amus* mutation, suggestive of normal HipHop functions despite defective splicing of its minSI. Therefore, the evolutionary advantage, if any, of creating a minSI for *hiphop* remains to be demonstrated.

### Rai1 is not essential for the elimination of unprotected U6-type snRNAs

We have shown that U6 and U6atac snRNAs both require an intact 5’ methyl-phosphate cap for stability (this study; Peng et al. 2023). We are interested in identifying the nuclease that carries out the degradation of uncapped snRNAs, and focused our attention on the DXO/Rai1 exonuclease. DXO/Rai1 has been shown to participate in the processing of RNAs with a variety of different 5’ structures and/or modifications (Doamekpor et al. 2019 and references therein). We reasoned that if Rai1 were responsible for degrading or initiating the degradation process of uncapped U6-type snRNAs, a double mutation of *rai1* and *amus* would restore to some extent the level of those snRNAs.

Using CRISPR/Cas9, we generated the *rai1^2-4^* allele with a 5bp deletion, creating a frameshift at codon 32 of 375 of the Rai1 protein (Figure 5A). Flies homozygous for *rai1^2-4^* or another *rai1* mutant allele *rai1^e04372^* are both viable and fertile. In addition, *rai1^2-4^; amus^1-1/A-1-3^* adults are phenotypically indistinguishable from those of *amus^1-1/A-1-3^*. As shown in Figure 5B, Northern blot analyses on total RNA isolated from adults did not reveal elevated levels of either U6 or U6atac under the double mutant background when compared to the *amus* single mutant. Therefore, we conclude that Rai1 is not responsible for degrading uncapped U6-type snRNAs.

**Figure 5.**
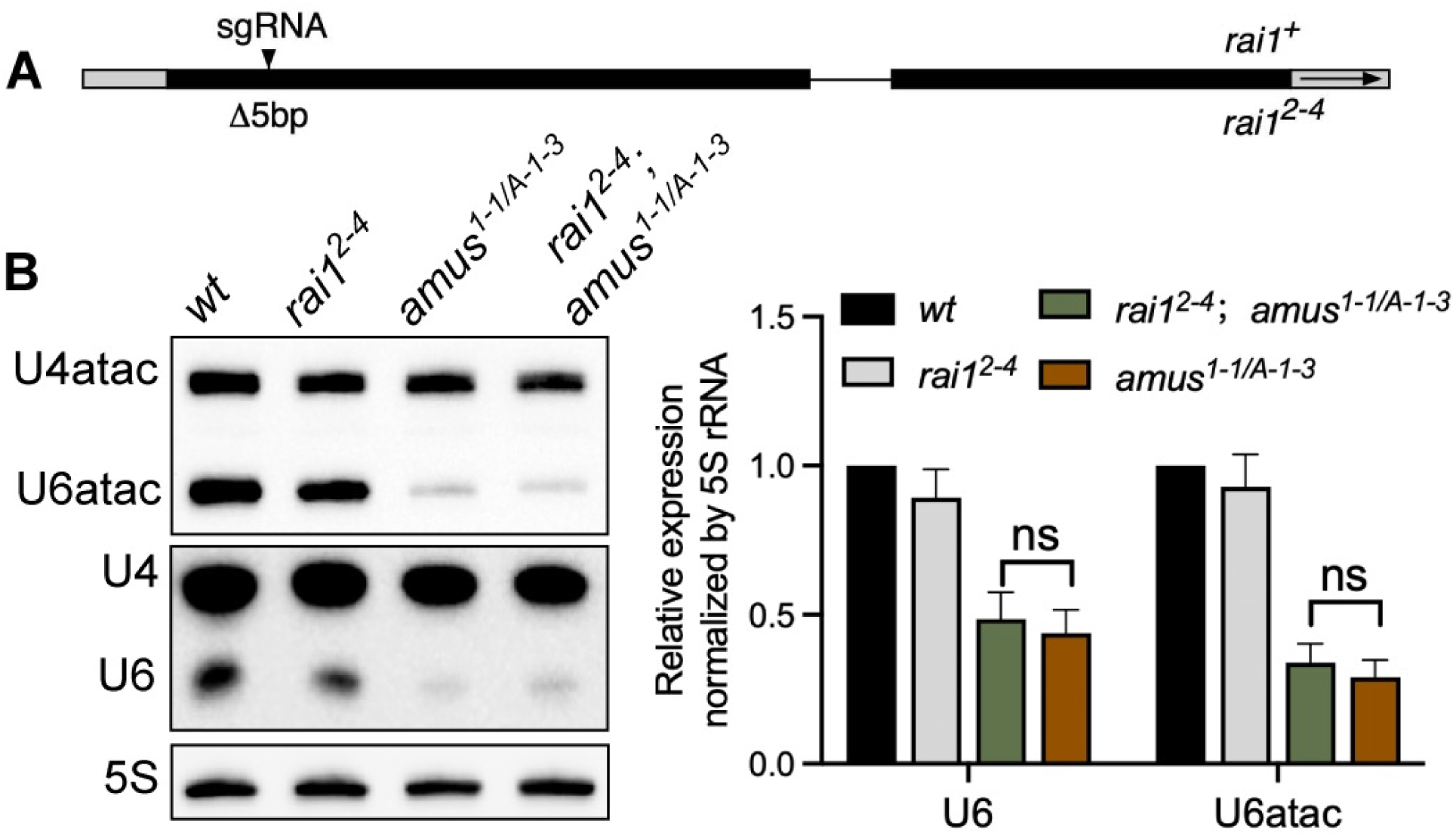
Rai1 is not required for the elimination of unprotected U6-type snRNAs. **A. Mutation of *rai1*.** A diagram of the *rai1* locus with black boxes representing coding region, grey boxes untranslated regions, and a horizontal line intron. At the top of the diagram, the position of the single sgRNA used in CRISPR/Cas9 mutagenesis is indicated, and 5bp deletion at the sgRNA targeted region of the *rai1^2-4^* allele is indicated at the bottom. **B. The effect of *rai1* mutation on small RNA production.** At the left is Northern blot analyses of small RNAs listed to the left of the pictures, with genotypes listed on top. RNA was isolated from adults. At the right is data quantification from three biological replicates. ns: not significant.

## Discussion

Here we showed that the U6-type snRNAs for both major and minor splicing are protected by the same enzyme. By leveraging a unique genetic background in which both major and minor splicing functions are compromised, we shed lights onto the regulation of a special class of introns that share features with both major and minor introns.

### “Minor spliceosome sensitive introns” are likely processed by both the major and minor spliceosomes

In previous works in which the “minor” function has been specifically disrupted, multiple approaches lead to evidence for the existence of a class of introns possessing “major” features that are processed instead by the minor spliceosome. That these introns are also under the control of the “major” function might be a foregone conclusion, but it is difficult to prove as total loss of major splicing has highly pleiotropic effects. Fortuitously, the *amus* hypomorphic background that we have generated partially disrupts both splicing functions by destabilizing the U6-type snRNAs. The genetic evidence supporting that our *amus* mutation is hypomorphic for both splicing functions is as followed. First, using the same RT-PCR based assay, we showed previously that intron retention is prevalent in major introns in *amus* mutant adults (Peng et al. 2023). Secondly, despite a greatly reduced U6atac level, the *amus* mutation does not abolish the splicing of canonical minor introns. However, a further reduction of the *U6atac* dosage enhances splicing defects to a level detectable by the RT-PCR assay. Genetically, this is consistent with a partial loss of minor function in the *amus* mutant. Despite the apparently normal splicing efficiency of the canonical minor introns, we clearly observed defects in minSI splicing while major introns in the same minSI-containing genes are not affected (Figure S2). The most parsimonious explanation, we suggest, is that the efficacy of minSI processing is already compromised in the sensitized background of U6atac reduction, and that the additional loss of the “major” function in *amus* mutants elevates the splicing defects to a detectable level.

Mechanistically, there might be some “leakiness” in splice site recognition so that minSIs can be “seen” by both classes of spliceosomes. A more provocative model for the existence of “hybrid” spliceosomes has been put forth that would contain a mixture of major and minor components, particularly involving the U4/U4atac and U6/U6atac pairs of snRNAs (Olthof et al. 2024). Experimentally, the production of chimeric snRNAs, by plasmid transfection into human cells, has proven that such a cross-talk between the two spliceosomes is not impossible (Shukla and Padgett 2004; Dietrich et al. 2009). Excitingly, a genetic test based on chromosomal copies of chimeric snRNAs is highly feasible in *Drosophila* as *U4atac* is a single copy gene, and only one of the three *U4* snRNA copies appear essential (Lu and Matera 2014).

### Minor-like introns are evolutionarily dynamic

Our limited sequence analyses nevertheless shed lights onto the evolution of the minor-like introns in *Drosophila*. We confirmed that these introns are generally less conserved than their canonical counterparts. Remarkably, almost half of the minSIs that we investigated display homologous splice site sequences only in the closest *Drosophila* species to *melanogaster*. Our RT-PCR validation of the *hiphop* intron appears consistent with this proposition. However, at least for the *hiphop* gene in *Drosophila*, the fact that a similarly positioned intron exists in some, but not other more distantly related species supports that the evolution of minSIs can also involve loss, in addition to interconversion between major and minor ones. Our result is also consistent with the suggestion that many of the introns in 5’UTR of human genes are likely evolving neutrally (Cenik et al. 2010).

### The evolutionarily conserved function(s) of the Amus/MePCE methyltransferases

The loss of the Amus methyltransferase destabilizes both U6 and U6atac snRNAs in *Drosophila*, which collectively control all pre-mRNA splicing in a cell. Therefore, loss of the mammalian MePCE protein is expected to impair splicing genome-wide, a functional consequence often ignored in prior studies of MePCE. MePCE is mostly studied for its function as an important transcriptional regulator *via* its role in maintaining the integrity of the 7SK snRNP (reviewed in Puidebat and Egloff 2025). MePCE not only caps, by methylation, but also binds to the 7SK snRNA, both contributing to the protection of 7SK. We and others have shown that the functions of mammalian MePCE as the capping enzyme for both 7SK and U6 have been divided between the Bin3 and Amus proteins in several animal species including *Drosophila* (Peng et al. 2023; Palumbo et al. 2024). Remarkably, Palumbo et al. (2024) showed that a Bin3 point mutation incapable of methylating RNA remains proficient in maintaining the stability of 7SK in *Drosophila*. This leads us to propose the following scenario to explain the evolutionary forces driving the Bin3 and Amus divergence. The ancestral Bin3/MePCE protein serves to methylate both 7SK and the U6-type snRNAs. For 7SK, it affords an additional mode of protection by RNA binding. After the ancestral split of MePCE into Bin3 and Amus, the enzymatic protection of U6-type snRNAs by Amus relieves the functional requirement of Bin3 as an enzyme without altering its ability to bind RNA. For the mammalian protein however, MePCE must remain as both an RNA binding protein and an active methyltransferase, as the loss of U6 or U6atac is lethal.

## Data Availability Statements

*Drosophila* strains, plasmids, and antibodies are available upon request. The authors affirm that all data necessary for confirming the conclusions of the article are present within the article, figures, and tables.

## Acknowledgements

We are grateful to the following scientists for sharing reagents: Dr. Yong-zhen Xu, Dr. Steven Hanes, Dr. Yirong Wang. We thank Dr. ShiShi Luo, Ms. Yiqing Wang for technical assistance.

---

**Figure S1.**
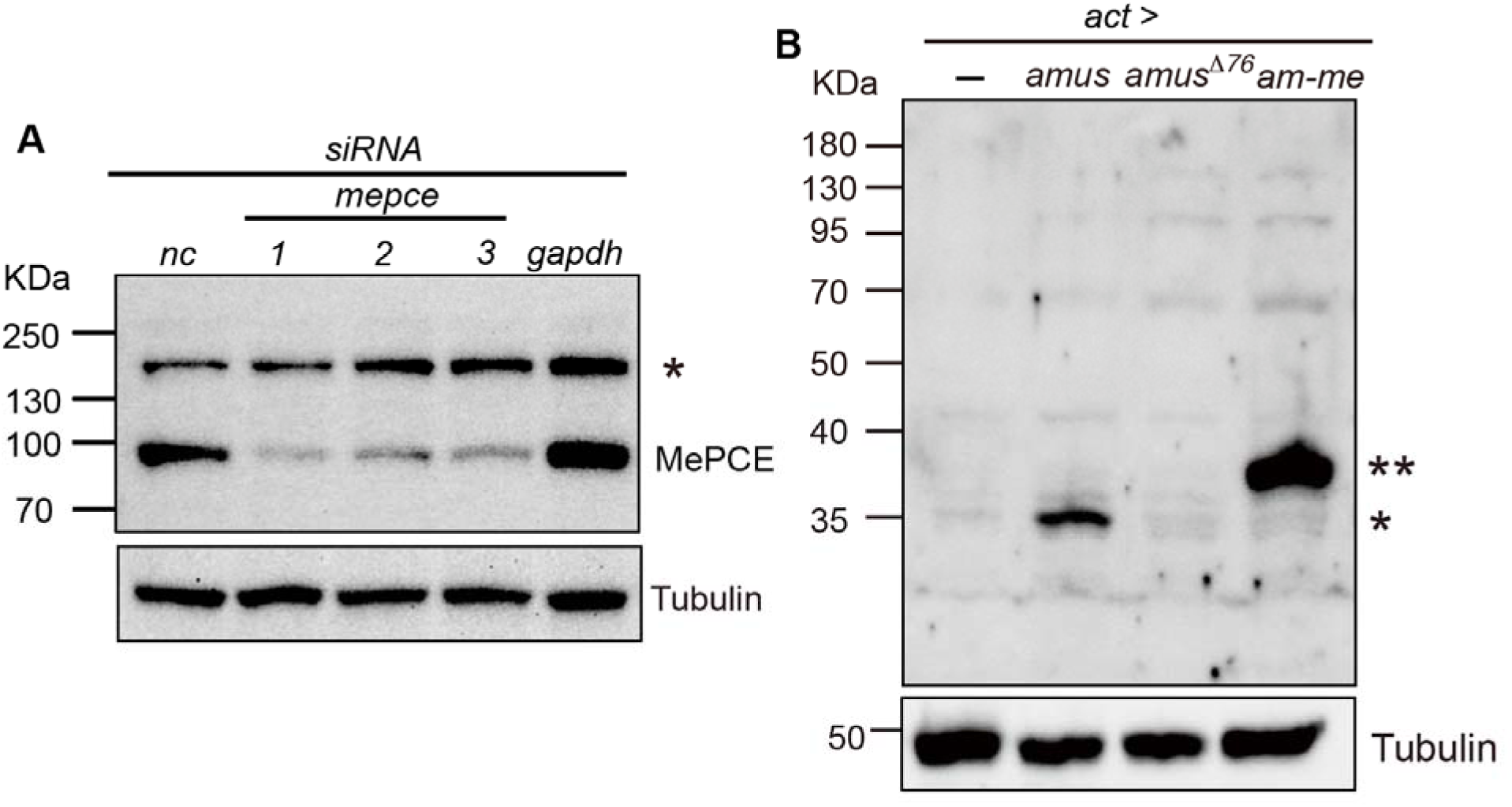
Monitoring protein levels of Amus and MePCE. **A. Western blot analyses of the effect of MePCE knock down.** Total extracts from HEK293T cells were loaded. On top of the panel showing Western blot results are RNAi treatment conditions. “nc” indicates a non-targeting siRNA as a negative control, with siRNA against *gapdh* as an additional control and three siRNA regents against different parts of *mepce* as previously described (Peng et al. 2023). The running positions of MePCE and the Tubulin control are indicated to the right, and the star indicates a non-specific band. **B. Western blot analyses of Amus overexpression in *Drosophila*.** Total pupal extracts with the indicated genotypes (top) were loaded. All four samples contain the *actin-gal4* transgene (act>). The leftmost sample contains no *amus* cDNA transgene (-), and is heterozygous for the *amus^A-1-3^* mutation. The other three samples were all homozygous for *amus^A-1-3^*, and contained: a cDNA for wildtype *amus* (amus); or a cDNA for Amus missing the first 76 residues (amus^Δ76^); or a cDNA for the Amus-MePCE chimeric protein (am-me). The star indicated running position of the full length Amus protein. The double star indicates running position of the Am-Me chimeric protein.

**Figure S2.**
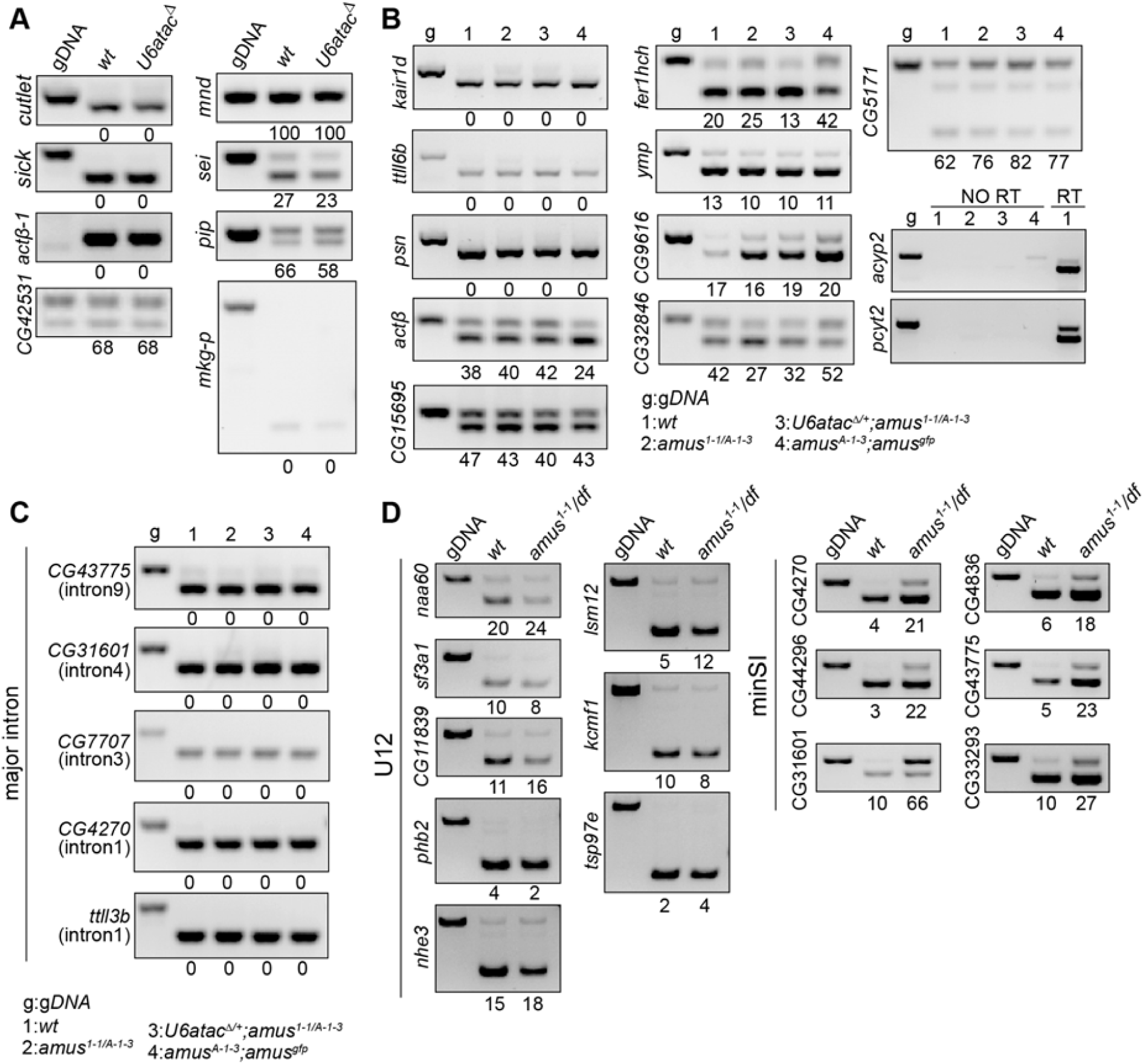
Additional RT-PCR assessment of minSI splicing. **A. Previously defined minSIs that are insensitive to the loss of U6atac.** RNA was isolated from larvae of the indicated genotypes. For the intron in the “*actbeta-1*” gene, the unspliced product is about 1700bp, therefore outside of the gel picture shown. **B. minSIs sensitive to the complete loss of U6atac but insensitive to the partial loss of Amus.** RNA was isolated from adults with the genotype indicated on top of the gel and listed at the bottom right. The bottom right panel shows RT-PCR results in which RT was omitted (NO RT) before PCR to assess the extent of genomic DNA contamination. The lanes with RT were from *wt* cDNA. **C. Major introns in the same genes as minSIs are not affected by *amus* mutations.** For each minSI-containing gene, a constitutive intron was chosen for the RT-PCR based splicing assay depicted in Figure 2A. RNA was isolated from adults with the genotype indicated on top of the gel and listed at the bottom. g: *wt* genomic DNA was used in PCR. **D. Testing the effect of another *amus* hypomorphic combination**. Examples are shown for the normal processing of canonical minor introns (U12), but abnormal processing of minSI introns in adults of *amus^1-1/df^*, which was heterozygous for a chromosomal deficiency of the *amus* region.

**Figure S3.**
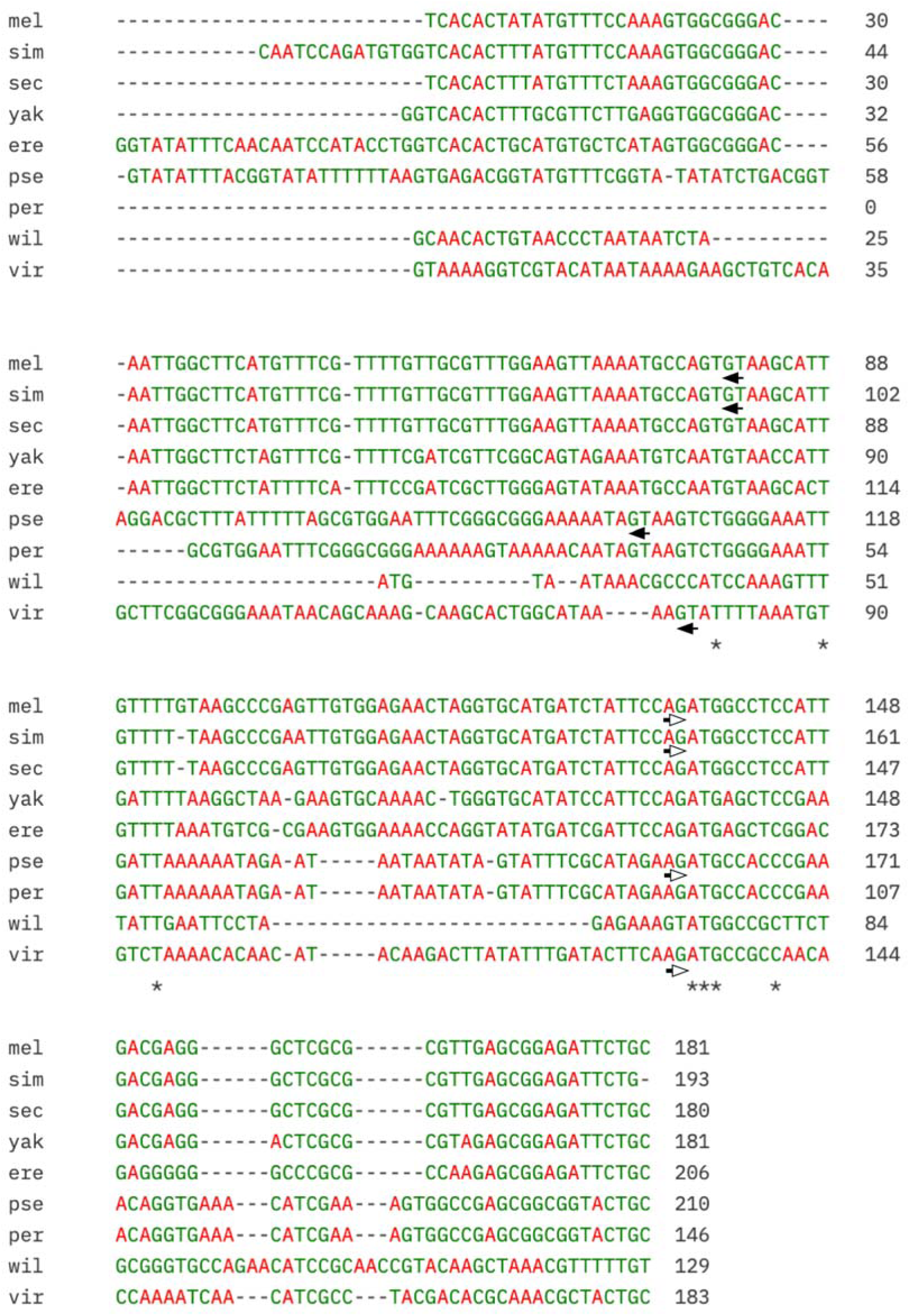
The intron in *hiphop* from *Drosophila* species. A Clustal alignment of the 5’ region of the *hiphop* genes from the following species: mel, *D. melanogaster*; sim, *D. simulans*; sec, *D. sechellia*; yak, *D. yakuba*; ere, *D. erecta*; pse, *D. pseudoobcura*; per, *D. persimilis*; wil, *D. willistoni*; vir, *D. virilis*. The 5’ boundary (left pointing solid arrow) and 3’ boundary (right pointing hallow arrow) of the introns are indicated for *D. mel*, *D. sim*, *D. pse*, and for *D. vir*. The ATG codon is next to the 3’ boundary.

**Figure S4.**
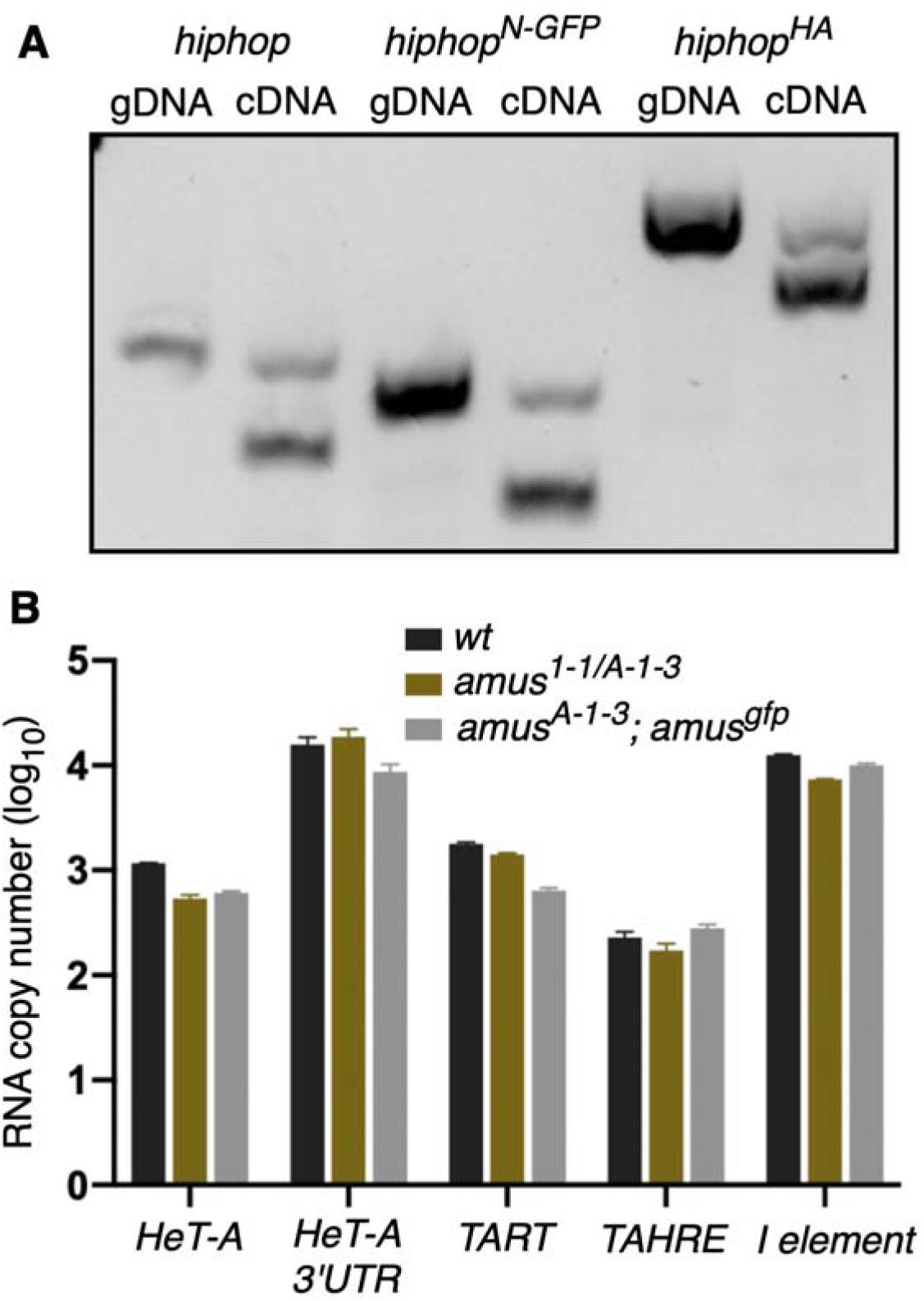
Functional characterization of the minSI of *hiphop*. **A. Sequence alterations at the 3’SS do not affect minSI splicing**. RT-PCR assay was performed using primers that flank the minSI and are specific to the *hiphop* alleles (shown on top). Genomic DNA (gDNA) was used as a template to mark the running position of the unspliced product in the reaction using cDNA as templates. **B. The *amus* mutation does not affect telomeric transposon expression**. qPCR quantification of telomeric expression was performed as described in Cui et al. 2021. Primers are specific to the *HeT-A* (two regions), *TART*, and *TAHRE* telomeric elements and the none telomeric *I element* as a control. Pair-wise comparisons show no significant difference between genotypes.

**Table S2.**
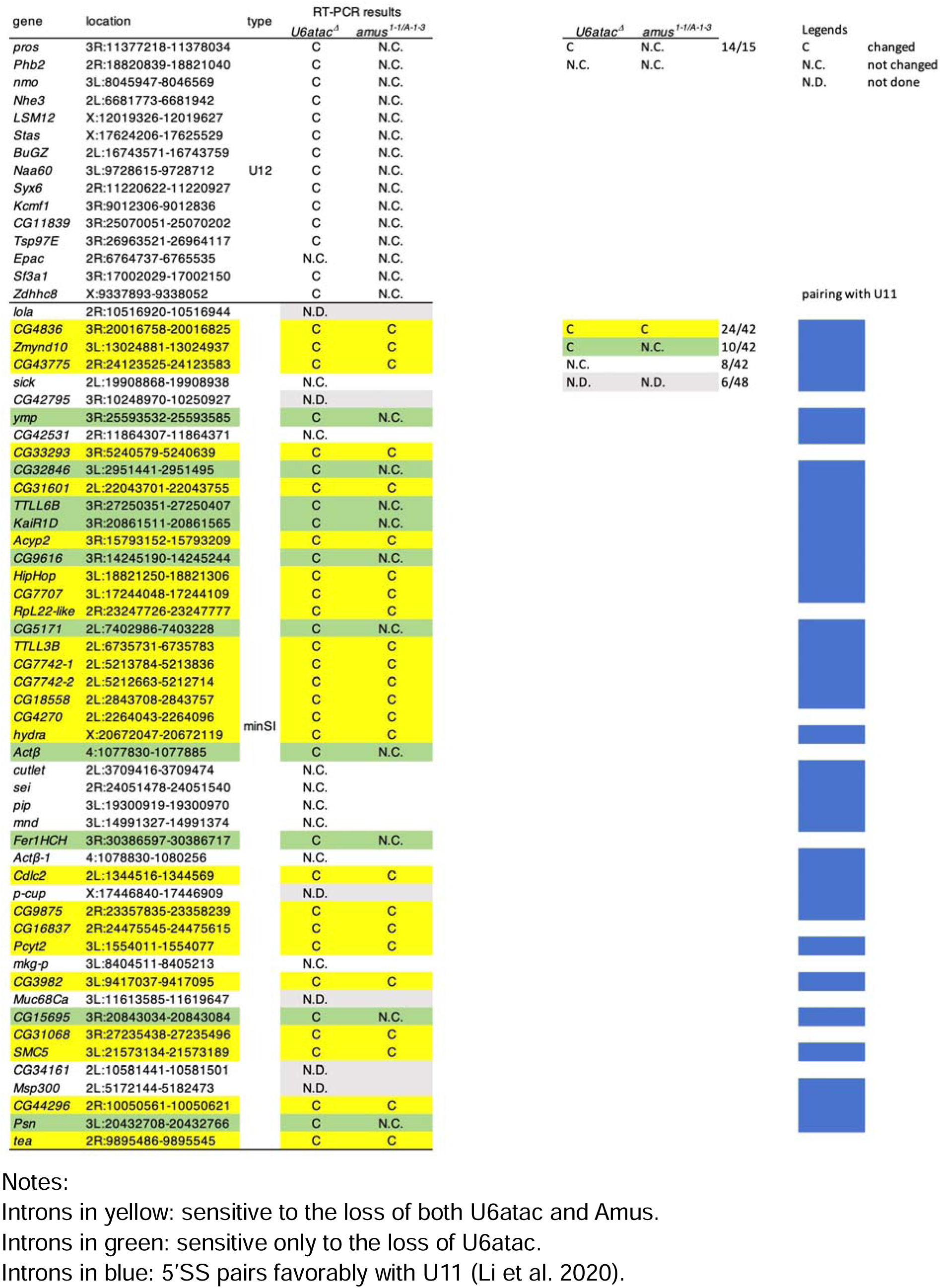
Summary of RT-PCR results of canonical (U12-type) and minS introns.

